# Sex differences in the epidemiology of tattoo skin disease in captive common bottlenose dolphins (*Tursiops truncatus*): are males more vulnerable than females?

**DOI:** 10.1101/101915

**Authors:** Marie-Françoise Van Bressem, Koen Van Waerebeek, Pádraig J. Duignan

**Author notes:** Corresponding author: M-F Van Bressem, Cetacean Conservation Medicine Group, Peruvian Centre for Cetacean Research, Lima 20, Peru.

## Abstract

The clinical and epidemiological features of tattoo skin disease (TSD), caused by cetacean poxviruses, are reported in 257 common bottlenose dolphins (*Tursiops truncatus*) held in 31 facilities in the USA and Europe. Photographs and biological data of 146 females and 111 males were analyzed. Dolphins were classified into three age classes (0-3; 4-8; over 9 years), approximating the life stages of ‘calves and young juveniles’, ‘juveniles and sub-adults’ and ‘adults’. The youngest dolphins with tattoos were 14 and 15 months old. Minimal TSD persistence varied between 4 and 65 months in 30 dolphins and was over 22 months in those with very large lesions (> 115 mm). In 2012-2014, 20.6% of the 257 dolphins had TSD. Prevalence varied between facilities from 5.6% (n= 18) to 60% (n= 20), possibly reflecting variation in environmental conditions. Prevalence was significantly higher in males (31.5%) than in females (12.3%), a pattern which departs from that observed in free-ranging Delphinidae where there is no gender bias. As with free-ranging Delphinidae, TSD prevalence in captive females varied with age category, being the highest in the 4 to 8 year old. By contrast, prevalence levels in males were high in all age classes. Prevalence of very large tattoos was also higher in males (28.6%, n= 35) than in females (11.1%, n= 18). Combined, these data suggest that captive male *T. truncatus* are more vulnerable to TSD than females possibly because of differences in immune response and because males may be more susceptible to captivity-related stress than females.

## INTRODUCTION

Captive common bottlenose dolphins (*Tursiops truncatus*) are affected by several cutaneous diseases, with the sub-acute form of erysipelas and tattoo skin disease (TSD) being the most notorious (Geraci et al. 1966, 1979, Sweeney and Ridgway 1975, Dunn et al. 2001). Erysipelas is caused by the pathogenic bacterium *Erysipelothrix rhusiopathiae* and, in the dermatologic form, is characterized by gray, elevated rhomboid lesions that occur over the entire body. The pathogenesis in this form presumably involves neutrophilic infiltration, bacterial microthrombosis of dermal vasculature and epidermal infarction leading to necrosis. Prompt antibiotic treatment is needed to avoid septicemia and death (Geraci et al. 1966, Sweeney and Ridgway 1975, Dunn et al. 2001). In odontocetes, TSD is characterized clinically by irregular, variably extensive, grey or black stippling of the skin (Geraci et al. 1979, Flom and Houk 1979, Van Bressem et al. 1993). Macroscopically, a presumptive diagnosis of TSD can be made based on visual inspection of high-resolution photography (Van Bressem et al. 2009). Histologically, there is focal vacuolar degeneration of cells in the *stratum intermedium* and compaction of adjacent cells in this layer. The overlying *stratum externum* is increased in depth by the proliferation of flattened cells extending into the epidermis rather than as an exophytic proliferation. The vacuolation of some epidermal cells and compaction and hyperplasia of others accounts for the gross appearance of pale foci with dark tattoo-like marks. Viral replication occurs in a transitional zone between the swollen vacuolated *stratum intermedium* and the compacted periphery as evidenced by eosinophilic cytoplasmic inclusion bodies visible on light microscopy and enveloped dumbbell-shaped virions on electron microscopy (Geraci et al. 1979, Flom and Houk 1979). In more chronic lesions, there is focal pitting and disruption of the surface layer allowing entry of bacteria and other opportunists (Geraci et al. 1979). The etiologic agents are poxviruses of the subfamily *Chordopoxvirinae,* tentatively classified into the genus *Cetaceanpoxvirus,* that is still awaiting acceptance (King et al. 2012). Cetacean poxviruses (CPVs) are genetically and antigenically more closely related to the orthopoxviruses than to the parapoxviruses (Bracht et al. 2006, Van Bressem et al. 2006, Blacklaws et al. 2013, Barnett et al. 2015, Fiorito et al. 2015). TSD is generally not a life-threatening disease (Geraci et al. 1979, Flom and Houk 1979, Smith et al. 1983). However, though its histological features and epidemiological pattern have been thoroughly investigated (Geraci et al. 1979, Flom and Houk 1979, Smith et al. 1983, Van Bressem and Van Waerebeek 1996, Van Bressem et al. 2003, 2009), its pathogenesis remains poorly understood.

In free-ranging Peruvian dusky dolphins (*Lagenorhynchus obscurus*), common dolphins (*Delphinus capensis* and *D. delphis*), *T. truncatus* and Hector’s dolphins (*Cephalorhynchus hectori hectori*) TSD prevalence levels varied significantly with age, being higher in juveniles compared to calves, attributed to the progressive loss of maternal immunity (Van Bressem and Van Waerebeek 1996, Van Bressem et al. 2009). Juveniles also had a higher probability of suffering TSD than adults, presumably because more adults had acquired active immunity following infection. A worldwide study of 1392 free-ranging odontocetes, comprising 17 species suggested that the epidemiological pattern and severity of TSD are indicators of population health (Van Bressem et al. 2009). In captive common bottlenose dolphins environmental conditions and general health may influence TSD expression, persistence and recurrence (Geraci et al. 1979, Smith et al. 1983).

Recent reports of severe and persistent TSD in captive dolphins in popular media prompted this investigation using methodology designed to document dermatitis in free-ranging dolphins (Van Bressem et al. 2003, Sanino et al. 2014). Here we report the clinical presentation and epidemiology of TSD in captive common bottlenose dophins at several facilities in the northern hemisphere.

## MATERIAL AND METHODS

### Dolphins

High quality images of 146 female and 111 male bottlenose dolphins held in 31 dolphinaria and marine parks in the USA (n= 18; F1-F18) and Europe (n= 13; F19-F31) from 2008 to 2014 were examined for the presence of TSD (Table 1). Images and biological data for each dolphin were initially retrieved from the Ceta-Base inventory (http://www.ceta-base.org), in agreement with the data holders. Additional photographs of the dolphins published on Flickr were also checked for the presence of tattoo skin lesions (often abbreviated as ‘tattoos’). Dolphins were classified into three age categories (0-3; 4-8; over 9 years), approximating the life stages of ‘calves and young juveniles’, ‘juveniles and sub-adults’ and ‘adults’, respectively (Read et al. 1993).

Codes used for facilities are: F1, Baltimore National Aquarium, MD; F2, Brookfield Zoo, IL; F3, Clear Water, FL; F4, Dolphin Cove/Dolphin Plus Bayside, FL; F5, Dolphin Plus/ Island dolphin care, FL; F6, Dolphin Research Center, FL; F7, Gulf World Marine Park, FL; F8, Gulfarium, Marine Adventure Park, FL; F9, Indianapolis Zoo, IN; F10, Institute for Marine Mammal Studies, MS; F11, Miami Seaquarium, FL; F12, Sea World San Antonio, TX; F13, Six Flags Discovery Kingdom, CA; F14, Texas State Aquarium, TX; F15, Discovery Cove, FL; F16 Mirage Dolphin Habitat, NV; F17, SeaWorld Orlando, FL; F18, SeaWorld San Diego, CA; F19, Acquario di Genova, Italy; F20, Boudewijn Park, Belgium; F21, Loro Parque, Spain; F22, Marineland Antibes, France; F23, OltreMare, Italy; F24, Selwo Marina, Spain; F25, Tampereen Sarkanniemi, Finland; F26, Tiergarten Nurnberg, Germany; F27, Zoo Aquarium de Madrid, Spain; F28, Parc Astérix, France; F29, Kolmardern Djupark and Zoo, Sweden; F30, Dolfinarium Harderwijk, The Netherlands; F31, Zoo Duisburg, Germany.

### Tattoo skin disease

Tattoos were identified on the basis of their macroscopic appearance, i.e. irregular, dark grey to black, cutaneous lesions with a stippled pattern. They were previously shown to be associated with poxviruses in free-living and captive *T. truncatus* by electron microscopy and/or polymerase chain reaction (Geraci et al. 1979, Flom and Houk, 1979, Van Bressem and Van Waerebeek 1996, Bracht et al. 2006). We calibrated lesion size relative to the maximum diameter of the dolphin’s eye (cornea) and height of its dorsal fin. ‘Small’ lesions (< 15 mm) corresponded to affected areas smaller than half of the corneal diameter; ‘medium-sized’ lesions (15-50 mm) had a diameter less than twice the corneal diameter, ‘large’ tattoos (51-115 mm) measured more than twice the corneal diameter and up to ½ dorsal fin height (Cartee et al. 1995, Sanino et al. 2014, Van Bressem et al. 2015). ‘Very large’ tattoos measured at least ½ dorsal fin height (> 115 mm). The age at first detection, the topography and the number of lesions were also examined. The minimal duration of the disease was examined using serial images of 30 positive dolphins taken in the period 2008 to 2014. We calculated TSD prevalence levels in the different facilities for the period 2012 to 2014 and examined the influence of sex and age categories on prevalence. Statistical significance of differences in prevalence (α = 0.05) was verified with chi-square tests.

## RESULTS

### Characteristics of TSD in captive dolphins

Tattoos were typical in all affected dolphins and were often associated with tooth rakes inflicted by conspecifics (Fig. 1). They were mostly distributed on the head, throat, flanks and back (Fig. 1) and ranged in number from one to more than 20 on the visible body parts. Hence these are minimum numbers. Tattoo size varied from small (5 mm) to very large (> 750 mm) (Fig. 1). The very large lesions were found in 10 males and two females ranging in age between 9 and 29 years (Fig. 2) and housed in the USA and in Europe (Table 1), respectively. Minimal TSD persistence varied between 4 and 65 months (mean= 32.8; SD= 21.0) in 30 dolphins and was over 22 months in those with very large lesions (Fig. 3). The youngest dolphins with TSD were 14 and 15 months old females born in the Netherlands in 2013. These calves had very few tattoos that varied in size from small to medium and in one calf they persisted for at least 4 months.

**Figure 1.**
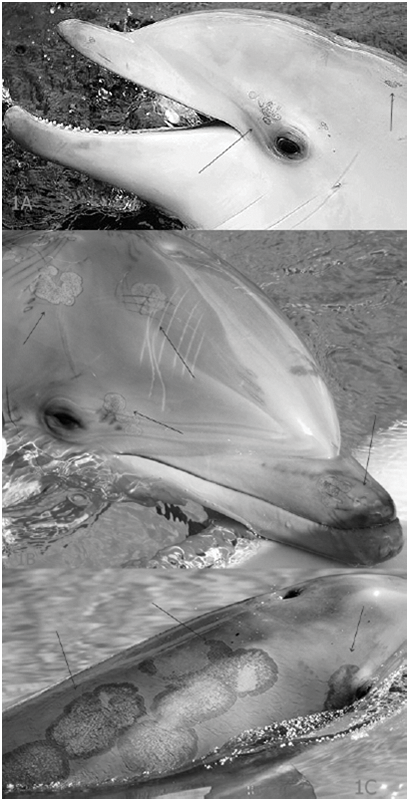
(a) Small and medium tattoos (arrows) with diagnostic stippled pattern on the head of a two year old male kept in captivity in The Netherlands, February 2014 © Robin de Vries. (b) Medium and large tattoos (arrows) on the head of a 5 year old male kept in San Diego, USA, May 2012. Various tattoos are topically associated with toothrakes (arrows) © Erin. (c) Very large tattoo lesions (arrows) on the back and head of an 11 year old male housed in Las Vegas, USA, August 2014 © Free the Mojave Dolphins.

**Figure 2.**
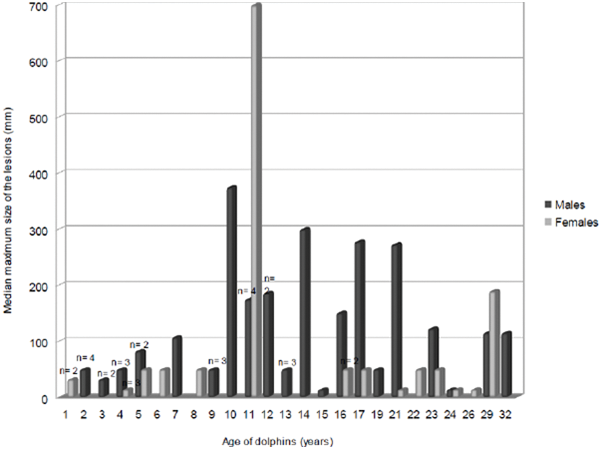
Variation in median maximum tattoo size (in mm) with age (in years) for 35 male and 18 female common bottlenose dolphins in facilities in the USA and Europe. Sample sizes for each age/sex class ranged n=1-4.

**Figure 3.**
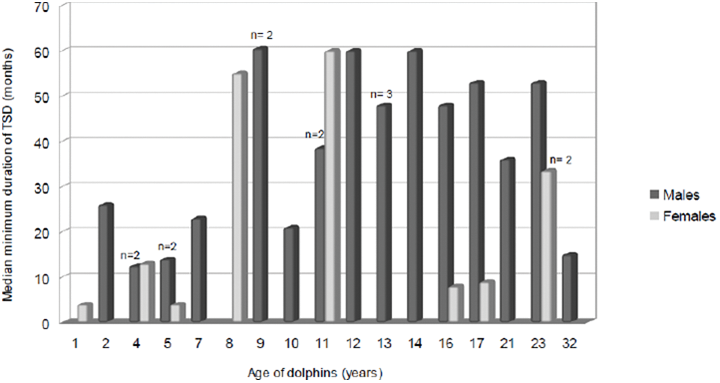
Median minimal duration of TSD in 30 captive bottlenose dolphins at facilities in the USA and Europe. Sample sizes for each age/sex ranged n=1-3.

**Table 1.** Prevalence (Prev) of tattoo skin disease in captive common bottlenose dolphin communities held in facilities in the Northern Hemisphere. Nt= total number of dolphins for which high quality images were available. Npos= number of dolphins positive for tattoo skin lesions. Affected facilities are indicated in italics. Symbol * indicates the presence of very large tattoo skin lesions in dolphins housed at these facilities. Code of each facility is provided in the Material and Methods.

| Facilities | Males |  | Females |  | All |  | Prev (%) |
| --- | --- | --- | --- | --- | --- | --- | --- |
|  | Nt | Npos | Nt | Npos | Nt | Npos |  |
| USA |  |  |  |  |  |  |  |
| F1 | 2 | 0 | 6 | 0 | 8 | 0 | 0 |
| F2 | 2 | 0 | 4 | 0 | 6 | 0 | 0 |
| F3 | 1 | 0 | 2 | 0 | 3 | 0 | 0 |
| F4 | 1 | 0 | 0 | 0 | 1 | 0 | 0 |
| F5 | 1 | 0 | 8 | 0 | 9 | 0 | 0 |
| F6 | 3 | 0 | 0 | 0 | 3 | 0 | 0 |
| F7 | 4 | 0 | 6 | 0 | 10 | 0 | 0 |
| F8 | 1 | 0 | 2 | 0 | 3 | 0 | 0 |
| F9 | 1 | 0 | 0 | 0 | 1 | 0 | 0 |
| F10 | 1 | 0 | 1 | 0 | 2 | 0 | 0 |
| F11 | 0 | 0 | 2 | 0 | 2 | 0 | 0 |
| F12 | 7 | 0 | 11 | 0 | 18 | 0 | 0 |
| *F13* | *10* | *1* | *8* | *0* | *18* | *1* | *5.6* |
| *F14\** | *2* | *1* | *0* | *0* | *2* | *1* | *50* |
| *F15* | *3* | *1* | *7* | *1* | *10* | *2* | *20* |
| *F16\** | *4* | *2* | *3* | *0* | *7* | *2* | *28.6* |
| *F17\** | *13* | *9* | *7* | *3* | *20* | *12* | *60* |
| *F18\** | *9* | *7* | *20* | *5* | *29* | *12* | *41.4* |
| Europe |  |  |  |  |  |  |  |
| F19 | 2 | 0 | 5 | 0 | 7 | 0 | 0 |
| F20 | 1 | 0 | 3 | 0 | 4 | 0 | 0 |
| F21 | 2 | 0 | 7 | 0 | 9 | 0 | 0 |
| F22 | 5 | 0 | 6 | 0 | 11 | 0 | 0 |
| F23 | 3 | 0 | 4 | 0 | 7 | 0 | 0 |
| F24 | 3 | 0 | 0 | 0 | 3 | 0 | 0 |
| F25 | 1 | 0 | 0 | 0 | 1 | 0 | 0 |
| F26 | 0 | 0 | 2 | 0 | 2 | 0 | 0 |
| F27 | 3 | 0 | 5 | 0 | 8 | 0 | 0 |
| *F28* | *5* | *2* | *4* | *0* | *9* | *2* | *22.2* |
| *F29\** | *0* | *0* | *3* | *1* | *3* | *1* | *33.3* |
| *F30* | *19* | *12* | *16* | *5* | *35* | *17* | *48.6* |
| *F31\** | *2* | *0* | *4* | *3* | *6* | *3* | *50* |
|  | 111 | 35 | 146 | 18 | 257 | 53 | 20.6 |

### Epidemiology of TSD in captive dolphins

In 2012-2014, 53 of 257 (20.6%) *T. truncatus* held in 31 marine parks in the USA and Europe had active TSD (Table 1). Six facilities were affected in the USA, and four in Europe (Table 1). Clinical TSD prevalence varied significantly (χ^2^= 15.05, df= 4, p= 0.005) between facilities with 10 or more dolphins, from 5.6% (n= 18, F13) up to 60% (n= 20, F17) (Table 1). TSD affected 18 females and 35 males (Table 1). Prevalence was significantly (χ^2^= 14.2, df= 1, p < 0.001) higher in males (31.5%, n= 111) than in females (12.3%, n= 146).

Tattoo prevalence varied with age categories in females, being the highest (23.1%) in the 4 to 8 year old (Fig. 4). However, this variation was not statistically significant (χ^2^= 3.4, df= 2, p = 0.18). In males, prevalence levels were high (above 25%) in all three age categories (Fig. 4).

Among affected dolphins prevalence of very large tattoos was higher in males (28.6%, n= 35) than in females (11.1%, n= 18), though not significantly so (χ^2^= 2.08, df= 1, p= 0.15). Prevalence of very large lesions varied in three facilities that housed more than 10 TSD-positive dolphins. It reached 50% in F17 (n= 12) and 16.7% in F18 (n= 12), but was zero in F30 (n= 17).

**Figure 4.**
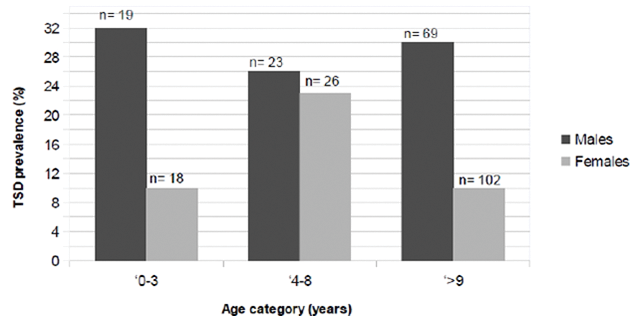
Variation in tattoo skin disease prevalence among age categories in male and female bottlenose dolphins at facilities in the USA and Europe. Age categories are: 0-3 years, 4-8 years, and older than 9 years.

## DISCUSSION

We applied techniques used to document TSD in free-ranging dolphins to study the epidemiology of infection in 257 captive common bottlenose dolphins in facilities in the Northern Hemisphere. This approach was previously validated by the authors by comparing electron microscopy and molecular diagnostic data with clinical presentation (Van Bressem et al. 1993, Van Bressem and Van Waerebeek 1996, Duignan 2000, Blacklaws et al. 2013). The disease was detected in 10 facilities in the USA and Europe with prevalence levels varying between 5.6% and 60% in the period 2012 to 2014 (Table 1). Differences in prevalence may reflect variation in environmental conditions, such as housing, water treatment methodology, crowding, noise, confinement, excessive or insufficient sun exposure and presence of aggressive individuals (Geraci et al. 1979, Dierauf 1990, Waples and Gales 2002, Fair et al. 2014) and/or the circulation of different strains of cetacean poxviruses with variable virulence in the affected facilities. By comparison, in three free-ranging *T. truncatu*s communities prevalence levels varied between 4.5% (Strait of Gibraltar, Spain, n= 334), 5.1% (Southeast Pacific Ocean, Peru, n= 79) and 21.9% (Sado Estuary, Portugal, n= 32) (Van Bressem et al. 2003, Jiménez-Torres et al. 2013). The dolphins residing in the Sado Estuary are regarded as a community under chronic physiological stress due to severe environmental degradation and high PCB exposure (Augusto et al. 2012, Jepson et al. 2016).

The earliest TSD infection among captive *T. truncatus* was detected in calves between 14 and 15 months old. Younger dolphins may still be protected by maternal immunity, as proposed to explain the significant age-related variation in prevalence encountered in several species of free-ranging odontocetes (Van Bressem and Van Waerebeek 1996, Van Bressem et al. 2009). Tattoo lesions were often associated with tooth rakes, suggesting that the virus could either be transmitted directly by bites, or that a damaged epidermis may facilitate infection. TSD persisted months and even years in some individuals and the lesions progressively grew to very large size. This pattern is consistent with cetacean poxviruses having developed immune evasion strategies, as reported for other poxviruses (Yousif and Al-Naeem 2012, Shisler 2015). This would favor viral persistence in marine parks. Further investigations on the pathogenesis of TSD in captive dolphins are clearly necessary.

Very large lesions were found in 10 males and two females between 9 and 29 year old, at six facilities (Table 1). In all cases the very large tattoos were observed in dolphins that had shown clinical TSD for more than 22 months. Prevalence of such unusual lesions was high among TSD positive dolphins kept at F17 (50%, n= 12) and F18 (16.7%, n= 12). Overall TSD prevalence was also high in these facilities (Table 1), suggesting environmental and/or social conditions that favor TSD occurrence and persistence. In free-ranging *T. truncatus* very large tattoos were observed only in a small community of dolphins in the Sado Estuary under chronic physiological stress (Van Bressem et al. 2003, Augusto et al. 2012, Jepson et al. 2016). Very large tattoos may be confused with infarctive epidermal necrosis caused by *E. rhusiopathiae* (Geraci et al. 1966, Dunn et al. 2001, Melero et al. 2011). However, the dermatitis of erysipelosis lacks the characteristic stippled pattern of tattoos, have a rhomboid, square or rectangular shape (cf. 'diamond skin disease') and run a very different clinical course, potentially leading to septicemia and death without antibiotic treatment (Dunn et al. 2001). Besides, erysipelosis is generally an historic disease of captive dolphins, since most facilities have eliminated clinical disease through vaccination (Lacave et al. 2001). TSD is a progressive, persistent disease in captive dolphins with no documented associated mortality, but also no reported treatment. There is no evidence of zoonotic transmission of TSD (Van Bressem et al. 2009). However, considering that several animal poxviruses can cause diseases in humans (Hicks and Worthy 1987, Haller et al. 2014), it would be sensible to take protective measures when handling TSD infected cetaceans and avoid all contact with delphinarium visitors.

In this study TSD prevalence was found to be 2.5 times higher in males than in females. This epidemiological pattern departs from that observed in free-ranging Delphinidae where there is no gender bias (Van Bressem and Van Waerebeek 1996, Van Bressem et al. 2009). In the captive females, prevalence varied with age category, being the highest in individuals between 4 and 8 years old. Similarly, in free-ranging Delphinidae the highest TSD prevalence levels occurred in juveniles (Van Bressem and Van Waerebeek 1996, Van Bressem et al. 2003, 2009). However, among males prevalence was high in all age classes. Males were also more likely to develop very large tattoo lesions than females though not significantly so. These data suggest that captive male *T. truncatus* are more vulnerable to TSD than females, possibly because of differences in immune response, because males are under greater social stress from conspecifics than females and because they may be more vulnerable to captivity-related stresses (husbandry, water quality, confinement). Excessive sun exposure has also long been recognized as an environmental stressor in captive dolphins (Dierauf 1990) and mitochondrial DNA damage associated with UV–induced microscopic lesions and apoptosis has recently been recently documented in three whale species (Bowman et al. 2013). Sexual maturation, reproduction and seasonal changes in testosterone can lead to increased stress levels in males through competition and aggression for access to females (Ridgway 1972, Connor and Smolker 1995, Waples and Gales 2002, Yamamoto et al. 2015). Numerous stress-related diseases, such as ulcerative gastritis, perforating ulcer, cardiogenic shock and psychogenic shock have been reported in captive cetaceans (Marino and Frohoff 2011). Interestingly, male *T. truncatus* kept in captivity in the USA and Portugal were also significantly more likely to be papillomavirus seropositive than females, while seroprevalence was not sex-dependent in dolphins free-ranging in Florida and South Carolina (Rehtanz et al. 2010).

Infection with cetacean poxvirus has long been recognized in facilities housing captive dolphins and there are many hypotheses regarding the transmission and maintenance of infection (Geraci et al. 1979, Flom and Houk 1979, Smith et al. 1983). Poxviruses likely circulate between facilities, as dolphins are regularly transferred from one aquarium to another. This procedure logically increases the risk of transmission between facilities and favors a broader geographic distribution of poxviruses and other pathogens in captive dolphins. Furthermore, shipment of dolphins between facilities is a very stressful event that may lead to alterations of the neuroendocrine response and to immune response impairment (Noda et al. 2007, Spoon and Romano 2012) and thus increase disease susceptibility. Combined, these data suggest that a high TSD prevalence together with a significantly higher occurrence in males than in females and the presence of very large tattoos, may be an indicator of chronic stress in captive bottlenose dolphins.

## CONCLUSIONS

1. Tattoo skin disease occurs at varying prevalence levels between facilities holding captive common bottlenose dolphin groups, the reason(s) for which should be further investigated.
2. TSD may persist for months and even years, and some dolphins, especially males, develop very large tattoos over time. The role of specific captive environmental conditions (husbandry, crowding, social structure, light intensity, etc.) should be studied as potentially contributing factors.
3. The cetacean poxviruses circulating in captive dolphins need to be characterized and compared with those infecting free-ranging populations. Though these viruses are not known to cause clinical disease in humans (Van Bressem et al. 2009), contacts between marine mammal park visitors and infected dolphins should be avoided.
4. Captive male *T. truncatus* have a significantly higher TSD prevalence than females with high prevalence levels in all age categories, and are more frequently affected by very large tattoos. Males may be more vulnerable to environmental and social stresses than females. Assessment of behavior, improvement of environmental quality and maintenance of appropriate groupings of age and sex classes should be attempted to reduce physiological and social stresses.

## Acknowledgements

We kindly thank Ceta-Base for giving us permission to use their database and providing additional information and contacts, Robin de Vries, ‘Free the Mojave Dolphin’ and Erin for allowing us to use their images and the many photographers on Flickr for posting photos with basic information on the dolphins. MFB thanks the Cetacean Society International (CSI) for sponsorship.

